# Deep-learning model–guided discovery and characterization of bacterial unspecific peroxygenases

**DOI:** 10.1101/2025.11.27.690923

**Authors:** Xiaowei Shen, Danning Song, Hui Zhang, Qian Yang, Biqiang Chen

## Abstract

Unspecific peroxygenases (UPOs) are capable of catalyzing the selective oxidation of organic substrates under mild conditions, using hydrogen peroxide (H_2_O_2_) as the sole oxidant. This makes them one of the most promising biocatalysts for chemical synthesis. However, the major limitation restricting the application of UPOs to date is their difficulty in heterologous expression. Although more than 4,000 putative UPO enzymes have been recorded in databases, only about 50 of them can currently be heterologously expressed. All UPOs discovered so far originate from eukaryotes (mainly basidiomycetes and some ascomycetes), and they rely on the complex expression systems and post-translational modifications of their native hosts. This further exacerbates the challenges associated with heterologous expression of eukaryotic UPOs. In this work, we developed a deep-learning-based enzyme mining strategy and, for the first time, discovered novel UPO enzymes from bacteria, achieving successful heterologous expression in Escherichia coli. Bacterial UPOs differ greatly from fungal UPOs in sequence similarity, displaying completely distinct evolutionary trajectories. The discovery of bacterial UPOs advances our understanding of the catalytic mechanisms and expression characteristics within the UPO family, breaking the long-held assumption that UPOs can only originate from eukaryotes. Their excellent heterologous expression performance and broad catalytic versatility will further expand the application potential of UPOs.

## Introduction

Selective oxyfunctionalization of unactivated C-H, C-C, and C=C bonds represents one of the most challenging tasks currently facing organic synthesis. Unspecific peroxygenases (UPOs) hold immense application potential as they can catalyze diverse reactions, such as epoxidation, hydroxylation, heteroatom oxidation, and halogenation, across a broad range of substrates under mild conditions using hydrogen peroxide (H_2_O_2_) as the sole oxidant^1,2^. Although cytochromes P450 monooxygenases catalyze the same oxyfunctionalization reactions as UPOs, UPOs function without the need for expensive cofactors (NAD(P)H) and auxiliary proteins. This renders them one of the most promising biocatalysts in chemical synthesis^2–5^, poised to dominate the new wave of oxyfunctionalization chemistry. Since the first UPO was discovered in *Agrocybe aegerita* in 2004^6^, there are now over 4,000 putative UPO enzymes present in current databases^7,8^. Phylogenetically, UPOs can be divided into two families with distinct structural and substrate characteristics: Family I (short UPOs) and Family II (long UPOs)^9^. Short UPOs are distributed across most fungal phyla and have an average size of 30 kDa. Long UPOs, on the other hand, have an average size of 44 kDa and have only been found in Ascomycota and Basidiomycota^10^. Long UPOs are monomeric proteins that possess an internal disulfide bridge and arginine as a charge stabilizer. Short UPOs are mostly dimeric, featuring an external disulfide bridge and histidine as a charge stabilizer^11^. Although both families display similar conserved structural and sequence motifs (the -EHD-S-E- and -EGD-S-R-E-motifs, as well as the typical PCP motif), they differ significantly in the organization (topology and size) of their heme access channel. This results in distinct substrate scopes: short UPOs accept larger substrates, whereas long UPOs show higher activity toward smaller compounds^12,13^. Despite the enormous potential of UPOs in biotechnology, the challenging heterologous expression in engineered strains remains a major technical bottleneck hindering their further large-scale application^10–12,14–17^. Although numerous studies have aimed at efficient expression in their natural hosts^18–21^, the results of homologous overexpression remain suboptimal due to the low genetic accessibility of fungal UPO strains and the limited availability of molecular tools for specific species. Consequently, the heterologous expression of UPOs in model organisms has become the primary research focus^10^. The first successful heterologous expression of a UPO occurred nearly a decade after the first UPO was discovered^22,23^, and currently, only around 50 enzymes are amenable to heterologous expression^14^.The strong dependence of these fungal UPOs on the complex expression systems and post-translational modifications (N- and/or C-terminal processing, S–S bridge formation, glycosylation, etc.) of their respective natural hosts^24^makes the heterologous expression of fungal UPOs exceptionally complex. Given the reality of difficult heterologous expression for fungal UPOs, the search for bacterial-derived UPOs appears to be a promising alternative, as prokaryotic expression systems are much simpler, rely less on complex post-translational modifications, and more readily enable high-level, soluble, and active expression at a lower construction and cultivation cost.

For a long time, UPOs were thought to exist only in fungi^15^; this was because UPOs require extensive and complex glycosylation modifications and the formation of correct disulfide bonds to exhibit catalytic activity, processes that are almost impossible to achieve in most prokaryotes. Extensive efforts in mining and identifying new UPOs have therefore focused exclusively on eukaryotes, neglecting the vast genetic resources within prokaryotes^25,26^. Traditional enzyme mining approaches heavily rely on sequence homology, and the enormous number of candidates leads to a heavy experimental verification burden. Consequently, only a tiny fraction can actually be expressed and assayed for activity, resulting in significant screening loss. Over the past five years or so, machine learning and deep learning have begun to fundamentally reshape enzyme mining itself. At the level of functional annotation, methods like CLEAN, which learns representations directly from raw amino acid sequences via contrastive learning-driven enzyme annotation, significantly outperform BLASTp in predicting EC numbers, particularly for enzymes with weak homology, scarce annotation, or multiple functions^27^. Structure-aware models like GraphEC further integrate predicted three-dimensional structures^28^with protein language model embeddings to simultaneously predict active sites, enzyme classification, and optimal pH within a graph neural network framework^29^. This makes it possible for researchers to *in silico* screen entire genomes or metagenomes for unannotated catalytic functions and environmentally adaptive enzymes before proceeding with experiments. With the growing understanding of UPO protein sequences, a large number of UPO-like sequences have been identified by searching databases of sequenced organisms. As whole-genome sequences become increasingly abundant, the number of annotated UPOs also increases. The current challenge is no longer the accessibility of gene sequences, but rather the accurate determination of sequence-function relationships and the selection of the most promising candidates from a vast pool of sequences for experimental verification.

Here, we introduce a new enzyme mining method, built upon our previous work, EITLEM-kinetics^30^, which can widely search existing protein sequence databases and rank candidate sequences based on their catalytic capability. This directly provides the most experimentally valuable enzymes, thus avoiding screening loss and alleviating the burden of extensive experimentation. We also report, for the first time, the discovery of UPO enzymes from bacteria. We successfully achieved the heterologous expression of these bacterial UPOs in *Escherichia coli* and conducted in-depth studies on their catalytic ability, substrate scope, and expression capacity.

## Results

### Construction of a novel enzyme mining method based on kinetic parameters prediction models

Enzyme kinetic parameters reflect the catalytic efficiency and substrate affinity of an enzyme, and the prediction of these parameters for protein-substrate pairs enables the mining of novel enzymes. Based on this, we developed a new enzyme mining method by integrating tools such as BLAST^31^and CD-HIT^32^into EITLEM-Kinetics **(Fig. 1)**. EITLEM-Kinetics achieves unified prediction of the enzyme’s *k*_*cat*,_ *K*_*m*_, and *k*_*cat*_/*K*_*m*_. Compared to similar models, EITLEM-Kinetics significantly improved prediction accuracy; for changes in catalytic capability resulting from amino acid mutations in the sequence, we tested the predictive power of EITLEM-Kinetics on mutants, achieving a coefficient of determination R^2^of 0.82 and a Spearman coefficient of 0.91. Regarding epistatic effects brought about by multiple mutations, we tested the model’s performance on data involving multiple point mutations and drastic fluctuations in kinetic parameters. The results showed that the model can still provide valuable predictions (with R^2^=0.663) even when the change magnitude exceeds 8284-fold and the number of mutation sites is as high as 18^30^Traditional enzyme mining methods based on sequence similarity struggle to quantify the enzyme’s catalytic capability; even within the same homologous family, the substrate scope, selectivity, and activity can vary widely, making differentiation based solely on sequence difficult. Because kinetic parameter prediction models inherently integrate substrate information for the reaction, and since kinetic parameters directly characterize enzyme catalytic efficiency and substrate affinity, this enables us to make a more targeted selection from massive homologous sequences.

**Figure 1.**
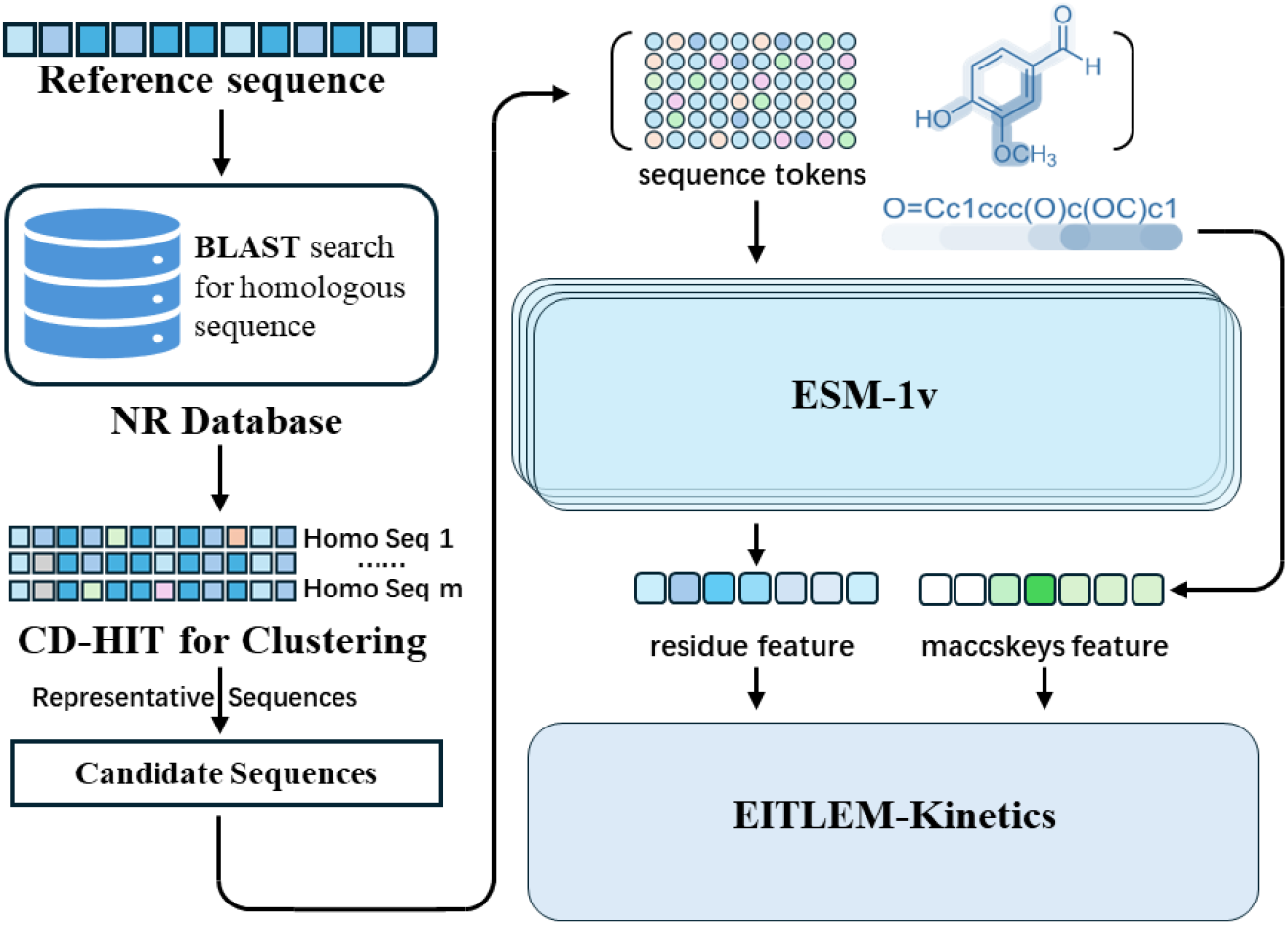
Architecture of the Novel Enzyme Mining Method Based on EITLEM-Kinetics. Homologous sequences are first searched in the non-redundant protein database (NR) using BLAST. The resulting homologous sequences are clustered using CD-HIT. EITLEM-Kinetics is then used to predict kinetic parameter values for the representative sequences, and the top N sequences with the highest predicted kinetic parameter values are selected for experimental verification.

### Mining of Bacterial-Derived UPOs

We performed mining in the non-redundant protein sequence database (NR database, as of December 18, 2024). In the first round, TteUPO was used as the reference sequence^12^, and NBD (3,4-Methylenedioxynitrobenzene, CAS 2620-44-2) was used as the model substrate to select the top 5 sequences ranked by predicted catalytic efficiency for experimental verification. In the second round, the highest-ranked sequence discovered in the first round, PseUPO, was used as the reference sequence, and the top 5 ranked sequences were selected for experimental verification **(Fig. 2a,b,c, in purple)**. The amino acid sequences of the bacterial UPOs are detailed in the Supporting Information.

**Figure 2.**
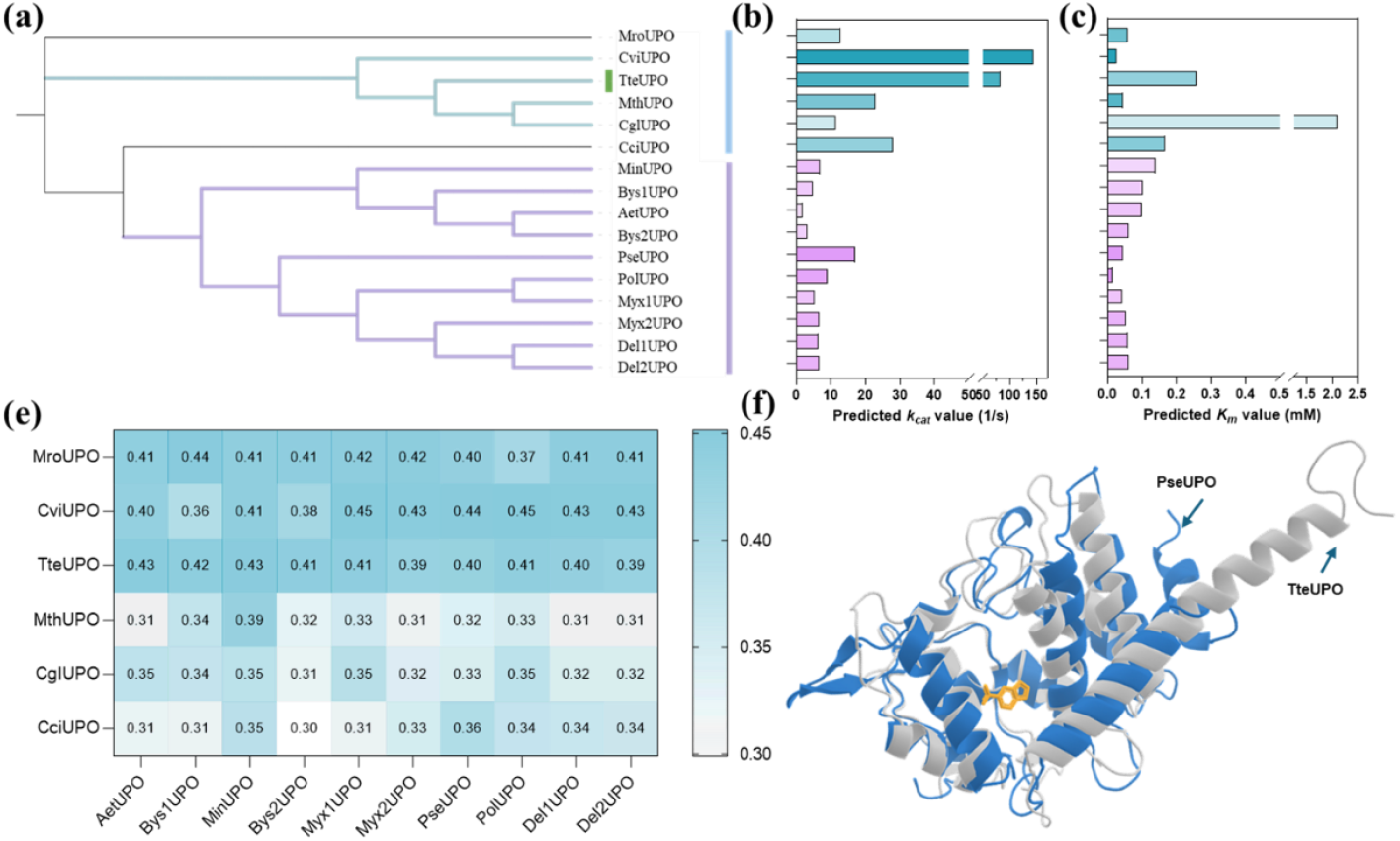
Comparison of Bacterial and Fungal UPOs. (a) Phylogenetic tree constructed for bacterial UPOs and six common fungal UPOs. (b) Predicted kcat values for fungal and bacterial UPOs. (c) Predicted Km values for fungal and bacterial UPOs. (e) Comparison of sequence similarity between bacterial UPOs and common fungal UPOs. (f) Structural comparison of the bacterial UPO PseUPO and the fungal UPO TteUPO.

Constructing a phylogenetic tree of bacterial UPOs and common fungal UPOs shows that fungal-derived nodes (e.g., MroUPO, CviUPO, etc.), colored blue, are mainly distributed on the left side or near the root region, while bacterial-derived nodes (e.g., MinUPO, Bys1UPO, etc.), colored purple, occupy the right branches and form multiple independent subclusters. Fungal and bacterial UPO sequences form two clearly distinct major branches. The fungal nodes are concentrated in a few branches, while the bacterial nodes are scattered and highly diverse, indicating a more significant evolutionary divergence for bacterial UPOs. The shorter branches of fungal UPOs suggest higher sequence conservation, which may be associated with specific oxidative metabolic functions they perform in fungi (such as lignin degradation, secondary metabolite synthesis), resulting in less evolutionary pressure. Bacterial UPOs are distributed across multiple dispersed branches (e.g., MinUPO, AetUPO, etc.), with considerable genetic distance between some subclusters, reflecting the rapid divergence of UPOs in bacteria driven by horizontal gene transfer (HGT) or environmental adaptation. The diversity of bacterial UPOs may be linked to their adaptive functions in different ecological niches (such as soil, host-associated environments). The diversity found in bacterial UPOs provides resources and possibilities for mining novel catalytic activities (such as substrate specificity).

Comparing the sequence similarity between bacterial UPOs and fungal UPOs **(Fig. 2e)** reveals a significant difference, with the maximum identity not exceeding 0.45. Modeling the reference sequence TteUPO and the highest-ranked predicted sequence, PseUPO, using AlphaFold3^33^**(Fig. 2f)** yielded a TM-score of 0.3834 and an RMSD of 3.48 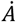. Generally, a TM-score > 0.5 is considered structurally similar, while a TM-score < 0.17 suggests random similarity. The value of 0.3834 indicates local structural conservatism but significant differences in overall folding. Although the overall sequence and folding differences are pronounced, locally conserved active sites may still retain function or even give rise to innovative catalytic centers. This implies that bacterial UPOs may possess catalytic capabilities for non-native substrates entirely different from those of fungal UPOs, warranting further exploration in the future.

### Expression of bacterial-derived unspecific peroxygenases (UPOs) in Escherichia coli

Given that all selected UPOs are of bacterial origin and transmembrane domain analysis **(see Supplementary Figure S4)** confirmed they are extracellular proteins, direct intracellular induced expression in Escherichia coli (E. coli) is prone to protein misfolding and inclusion body formation. The E. coli auto-induction expression system enables slow synthesis of target proteins through metabolic regulation of nutrients, which not only significantly improves the intracellular soluble expression efficiency of extracellular proteins but also supports high-density growth of E. coli via the abundant nutrients in the medium, with the bacterial culture OD600 value reaching 10. Therefore, the auto-induction system was preferentially used for the heterologous expression of UPOs in this study. The final expression levels of each strain are as follows **(Fig. 3)**: Myx1UPO (28.2 mg/L), Myx2UPO (30.0 mg/L), PseUPO (34.0 mg/L), Del1UPO (32.0 mg/L), Del2UPO (33.0 mg/L), and PolUPO (35.0 mg/L). Among them, PolUPO exhibited the highest expression level, while Myx1UPO showed the lowest. The expression levels of all strains ranged from 28.2 to 35.0 mg/L, demonstrating excellent expression stability.

**Figure 3.**
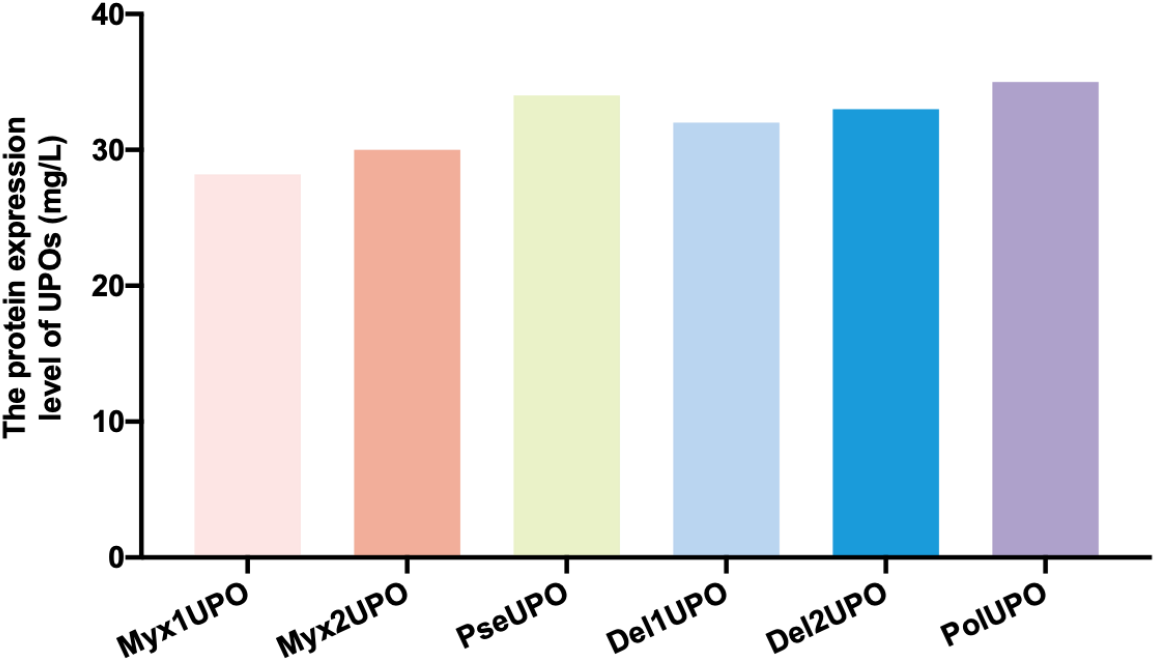
Expression levels of bacterial-derived unspecific peroxygenases (UPOs) (mg/L)

### Enzymatic characterization of bacterial-derived unspecific peroxygenases (UPOs) towards aromatic substrates

All bacterial-derived unspecific peroxygenases (UPOs) selected in this study exhibited catalytic activity towards three substrates, namely 1,2-methylenedioxy-4-nitrobenzene (NBD), veratryl alcohol, and naphthalene, with substrate-specific reaction types: dealkylation of NBD to catechol, hydroxyl oxidation of veratryl alcohol to veratraldehyde, and hydroxylation of naphthalene to 1-naphthol. Significant differences were observed in the catalytic activity (expressed as U/L) and catalytic efficiency (expressed as *k*_*cat*_*/K*_*m*_, unit: mM^−1^·s^−1^) of each UPO towards different substrates **(Fig 4)**.

**Figure 4.**
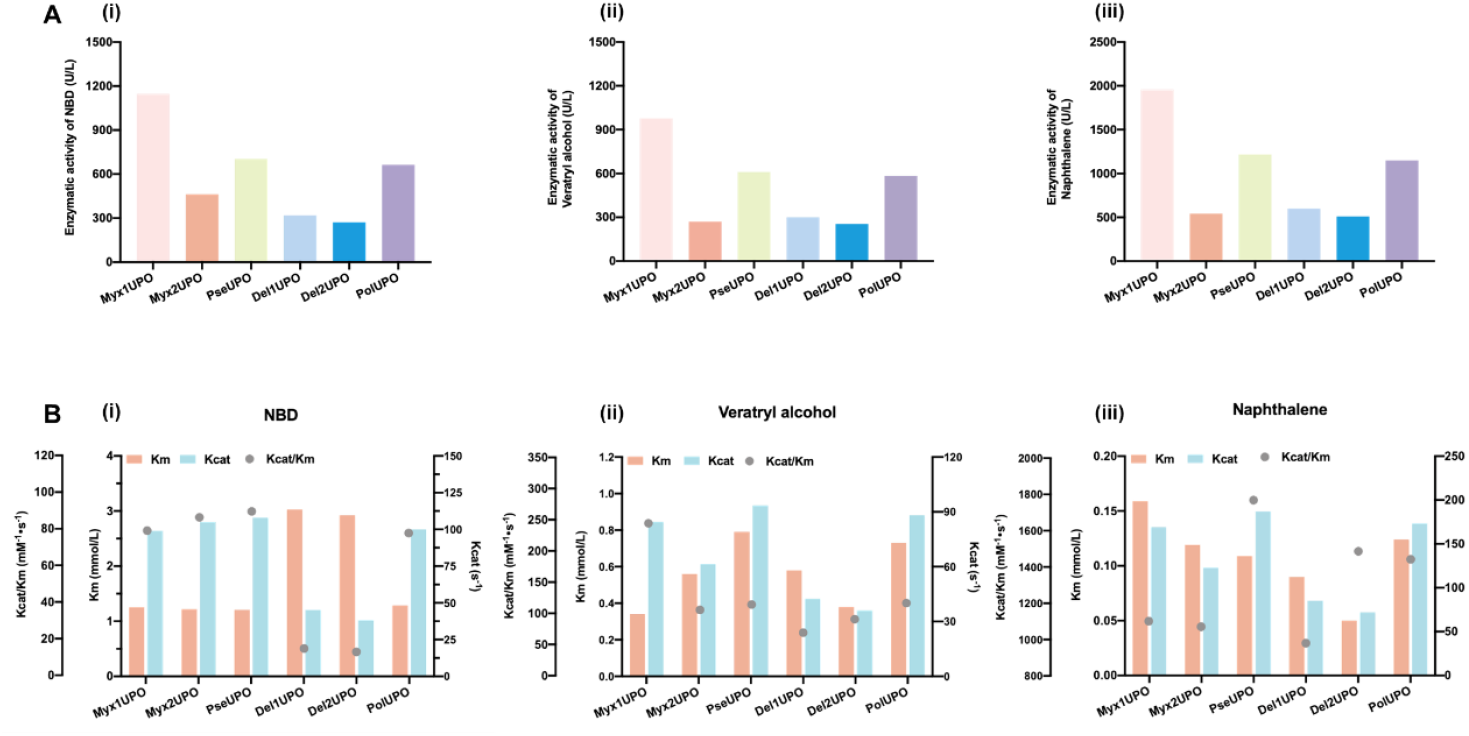
Catalytic activities (U/L) and catalytic efficiencies *k*_*cat*_*/K*_*m*_, mM^−1^·s^−1^) of UPOs towards aromatic substrates: (A) Catalytic activity; (B) Catalytic efficiency.

All UPOs displayed significantly higher catalytic activity towards naphthalene (510.04-1959.00 U/L) than towards NBD and veratryl alcohol, with the highest catalytic efficiency (kcat/Km = 944.89-1716.79 mM^−1^·s^−1^), indicating that naphthalene is the preferred substrate for this class of UPOs. The catalytic activity towards veratryl alcohol was generally the lowest (254.95-978.00 U/L), but the variation in catalytic efficiency was relatively small (73.28-248.59 mM^−1^·s^−1^).

The catalytic activities and efficiencies (*k*_*cat*_/*K*_*m*_) of the UPOs against three model substrates (NBD, veratryl alcohol, and naphthalene) are shown in Figure 4. Myx1UPO demonstrated the highest catalytic activity across all three substrates, notably reaching 1959.00 U/L for naphthalene, generally followed by PseUPO and PolUPO. The ranking of catalytic efficiency, however, showed substrate dependency: PseUPO exhibited the highest *k*_*cat*_/*K*_*m*_ for NBD (89.40 mM^−1^·s^−1^), Myx1UPO showed the highest efficiency for veratryl alcohol (248.59 mM^−1^·s^−1^), and PseUPO again topped the ranking for naphthalene (1716.79 mM^−1^·s^−1^), with Del2UPO and PolUPO also showing high activities.

In conclusion, Myx1UPO exhibited the highest catalytic activity towards all three substrates, especially for naphthalene (1959.00 U/L), which was significantly superior to other strains. PseUPO showed the optimal comprehensive catalytic efficiency, with the highest *k*_*cat*_/*K*_*m*_ value for naphthalene (1716.79 mM^−1^·s^−1^) and leading catalytic efficiency for NBD and veratryl alcohol. Del1UPO and Del2UPO displayed relatively low catalytic activity and efficiency towards NBD, but Del2UPO exhibited excellent catalytic efficiency for naphthalene.

### Catalytic functions of bacterial-derived unspecific peroxygenases (UPOs) towards terpenoid substrate

The screened bacterial unspecific peroxygenases were all capable of catalyzing the transformation of three terpene substrates—limonene, pinene, and 3-carene—with reactions primarily involving epoxidation, hydroxylation, and carbonylation. However, significant enzyme-specific differences in chemoselectivity were observed across these substrates **(Fig. 5)**.

**Figure 5.**
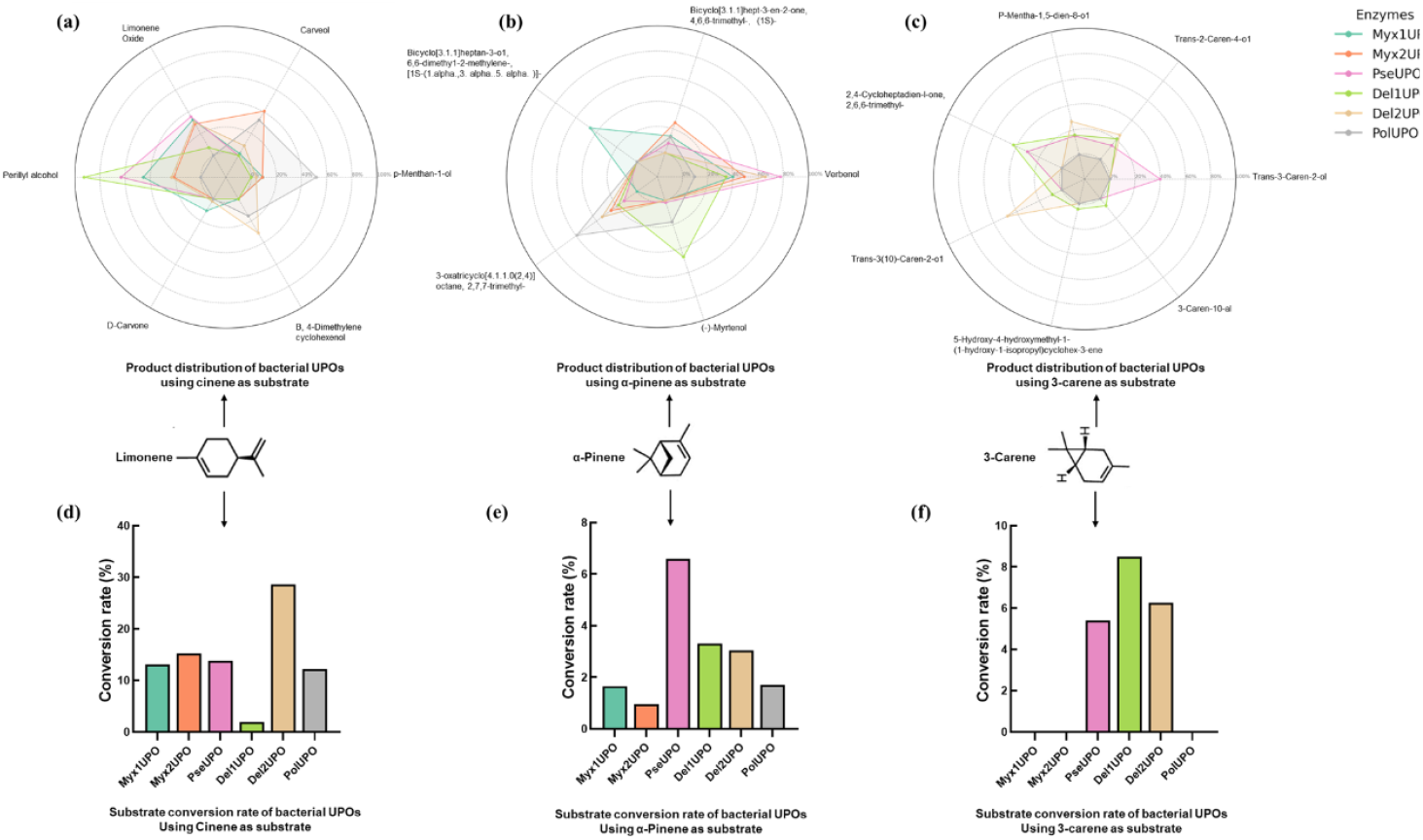
Multi-substrate catalytic experiments of bacterial UPO. (a)-(c): The product distribution of bacterial UPO with Cinene, α-Pinene and 3-Carene as substrates, respectively; (d)-(e): The conversion rate distribution of bacterial UPO with Cinene, α-Pinene and 3-Carene as substrates, respectively.

For limonene **(Fig.5a,d)**, all UPOs were effective catalysts. Myx1UPO and PseUPO showed high preference for allylic methyl hydroxylation (yielding perillyl alcohol, up to 92.74% for Del1UPO) and double-bond epoxidation (limonene oxide), with Myx1UPO exhibiting the highest overall activity. Del1UPO was notable for its exceptional selectivity toward methyl hydroxylation (92.74%), while Myx2UPO and PolUPO showed broader activity profiles, including ring hydroxylation and subsequent oxidation to D-carvone or cyclohexanol (PolUPO leading in cyclohexanol formation at 51.82%). Del2UPO demonstrated the most diverse product profile, including the unique beta-menthan-1-ol product.

For pinene **(Fig.5d,4)**, all UPOs were active, with the core reaction types being ring hydroxylation, double-bond epoxidation, and hydroxyl oxidation/carbonylation. Myx1UPO exhibited extremely high ring hydroxylation selectivity (86%, yielding verbenol and its isomers), with partial subsequent oxidation. PseUPO similarly favored ring hydroxylation (77.83%). In contrast, PolUPO primarily favored double-bond epoxidation (59.29%). Del1UPO was distinctive for having the highest selectivity toward methyl hydroxylation (myrtenol, 47.1%), whereas Myx2UPO showed a balanced presence of all three core reaction types.

For 3-carene **(Fig.5e,f)**, only PseUPO, Del1UPO, and Del2UPO} were effective. Their selectivity profiles were distinct: Del2UPO exhibited absolute hydroxylation selectivity (100%). PseUPO mainly catalyzed ring methyl alpha-hydroxylation and prop-methyl hydroxylation, with a notable 30.52% of the product resulting from allyl oxidation to ketones. Del1UPO displayed the highest oxidizing power among the active enzymes, with its initial hydroxylation products further oxidized to ketones (42.9% selectivity), combining both hydroxylation (49.9%) and subsequent oxidation activity.

## Conclusion

In this work, we successfully pioneered a deep-learning-guided enzyme mining strategy built upon our EITLEM-Kinetics model, which efficiently predicts and ranks enzyme candidates based on their kinetic parameters (*k*_*cat*,_ *K*_*m*_, and *k*_*cat*_/*K*_*m*_). This approach allowed us to move beyond traditional homology-based screening, targeting enzymes with high intrinsic catalytic potential and alleviating the burden of extensive experimental verification. Most significantly, our methodology led to the unprecedented discovery of unspecific peroxygenases (UPOs) from bacterial sources, a finding that definitively overturns the long-standing paradigm limiting UPOs to eukaryotic organisms, primarily fungi. The newly discovered bacterial UPOs are evolutionarily distinct, showing low sequence identity and forming separate phylogenetic branches from their fungal counterparts. This genetic and potential structural divergence is highly valuable. Crucially, we demonstrated that these bacterial UPOs can be successfully and robustly expressed in the simple prokaryotic host Escherichia coli at high titers (up to 35.0 mg/L) using an auto-induction system. This achievement directly addresses the major technical hurdle of difficult heterologous expression that has plagued the widespread application of fungal UPOs, thus making large-scale production a far more accessible reality. Our comprehensive characterization confirmed the broad and versatile catalytic power of these enzymes. They efficiently catalyzed the selective oxidation of various substrates, including aromatic compounds (dealkylation, hydroxylation) and complex terpenes (epoxidation, hydroxylation, carbonylation). Enzymes such as Myx1UPO and PseUPO demonstrated superior activity and catalytic efficiency across different model substrates, while others, like Del1UPO, exhibited unique chemoselectivity profiles, particularly in terpenoid transformations. In summary, the discovery of a new, highly-expressible family of UPOs in bacteria significantly expands the biocatalytic toolkit for selective oxyfunctionalization, providing robust and cost-effective alternatives for chemical synthesis and opening new avenues for understanding the evolutionary history and potential of the UPO enzyme superfamily.

## Author contributions

X.S. was responsible for conceptualizing the project, developing the methodology, ensuring validation of the work, creating and maintaining the software. D.S. was primarily responsible for experimental operations, sample preparation, and data validation. H.Z. was responsible for the implementation of the experimental design and data collection, and provided support in the initial drafting of the manuscript. B.C. and Q.Y. played a key role in conceptualization and methodology development, was involved in the review and editing of the written content.

## Additional information

### Supplementary information

The online version contains supplementary material available.

