## Supporting Information for "Deep-learning model–guided discovery and characterization of bacterial unspecific peroxygenases"

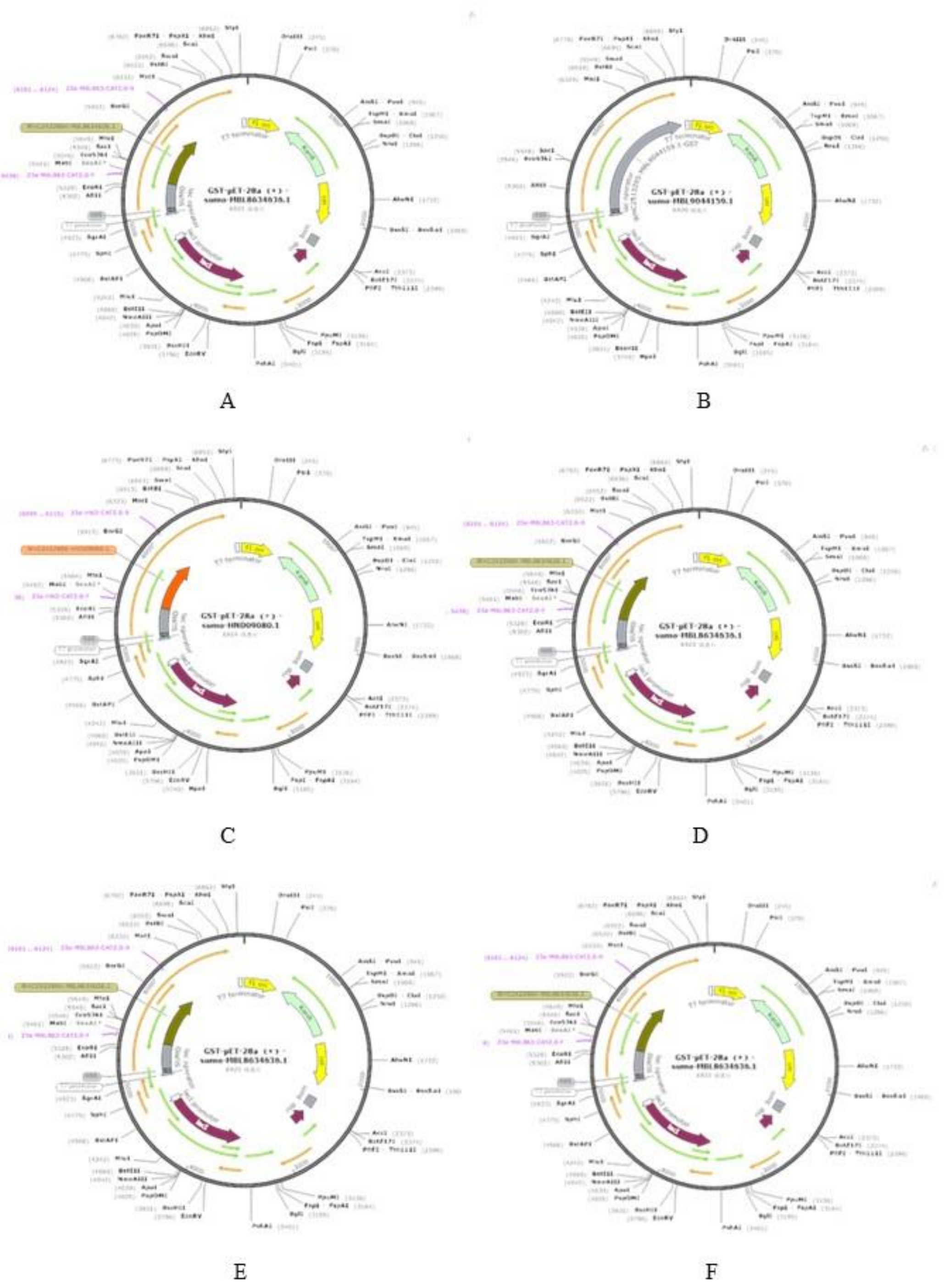


**Figure S1.** Plasmid maps A:pET28a-SUMO-Myx1UPO-GST，B:pET28a-SUMO- Myx2UPO-GST，C: pET28a-SUMO- PseUPO-GST，D: 28a-Del1UPO-GST，E: 28a-Del2UPO-GST，F: 28a-PolUPO-GST

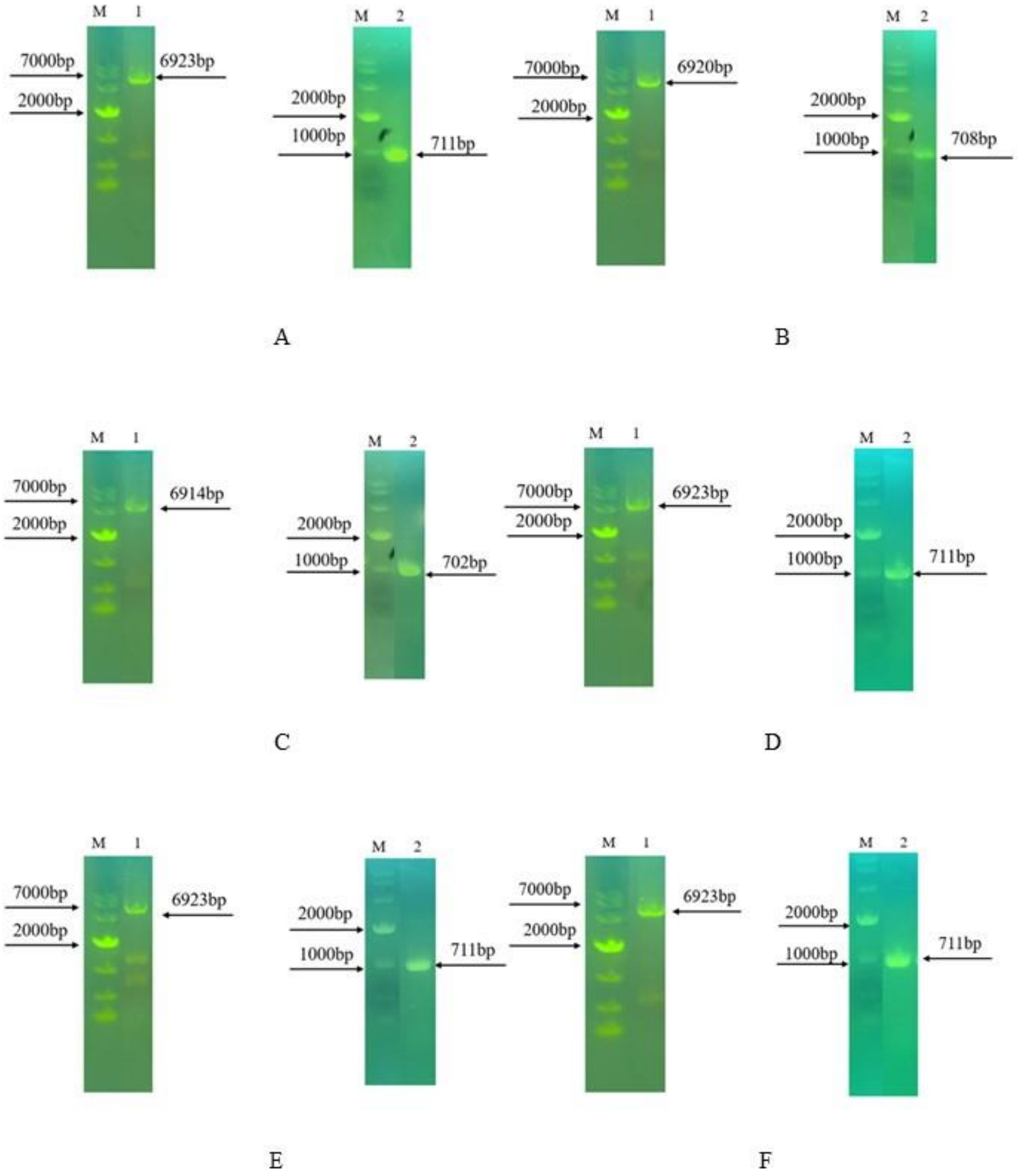


**Figure S2.** DNA agarose gel electrophoresis profiles，M：10000bp 的 Marker；A:Myx1UPO、B:Myx2UPO、C:PseUPO、D:Del1UPO、E:Del2UPO、F:PolUPO

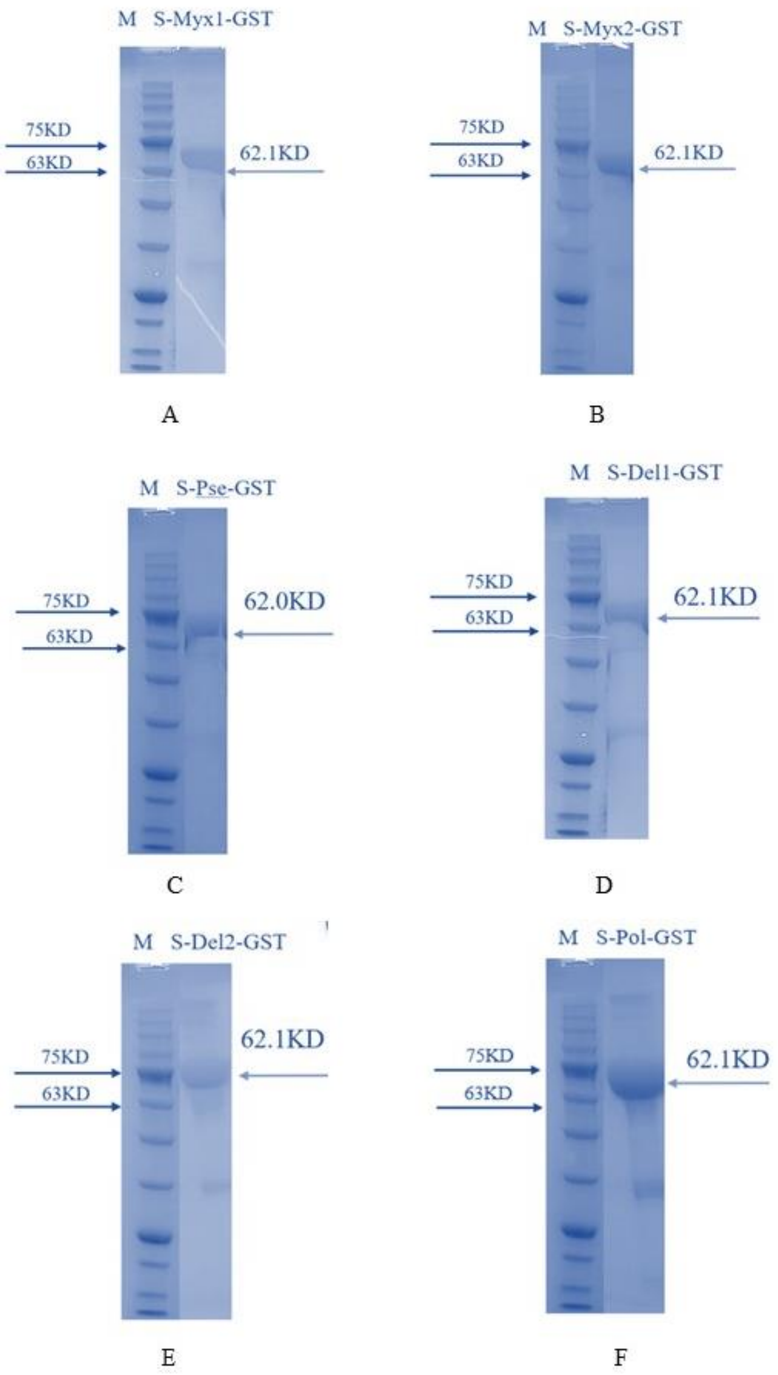


**Figure S3.** Protein purification profiles A:Myx1UPO、B:Myx2UPO、C:PseUPO、D:Del1UPO、E:Del2UPO、F:PolUPO

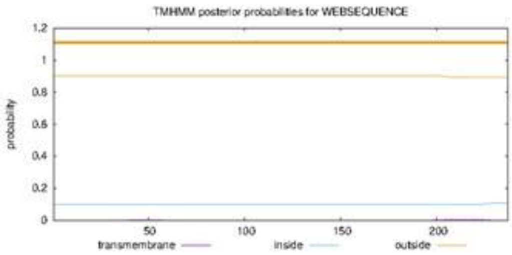


Myx1UPO

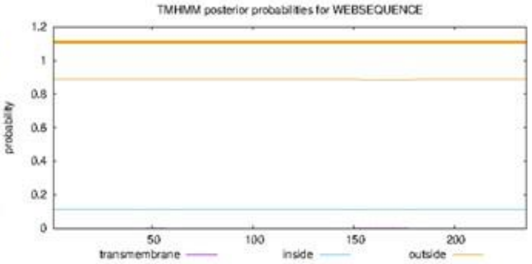


Myx2UPO

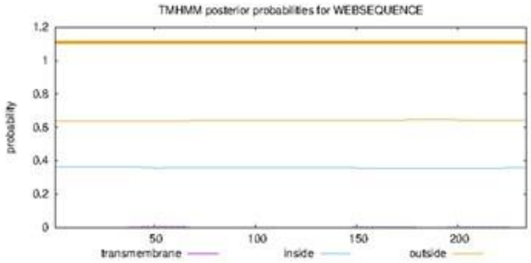


PseUPO

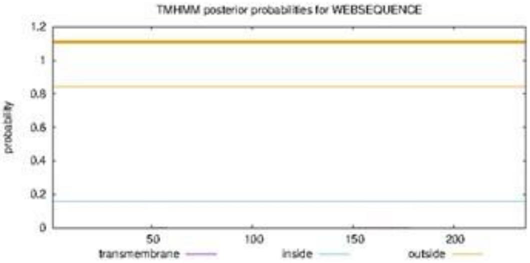


Del1UPO

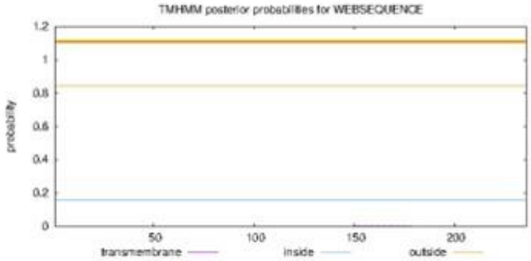


Del2UPO

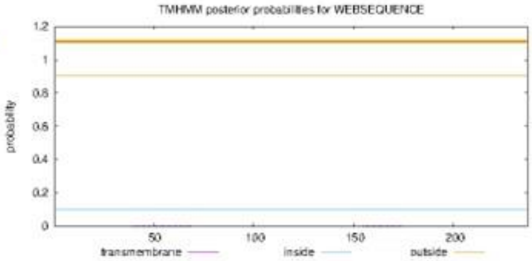


PolUPO

**Figure S4.** Transmembrane domain analysis

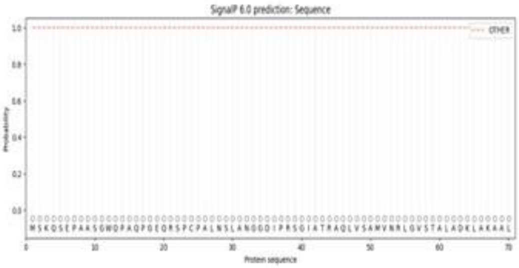


Myx1UPO

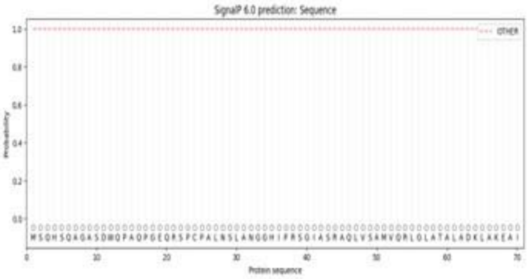


Myx2UPO

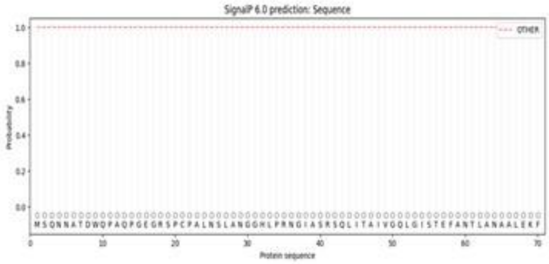


PseUPO

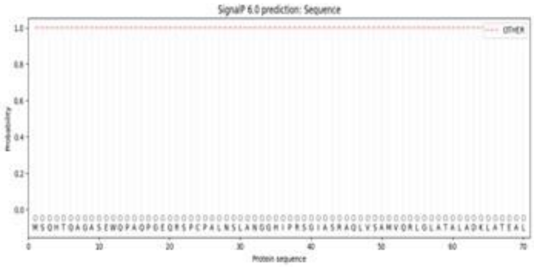


Del1UPO

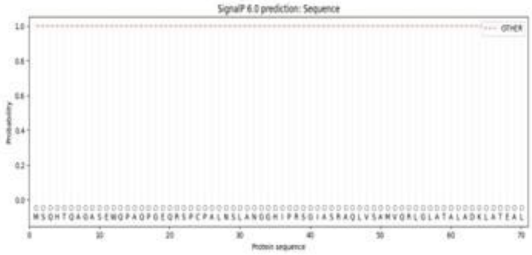


Del2UPO

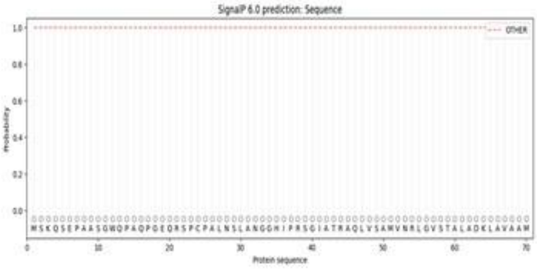


PolUPO

**Figure S5.** Signal peptide prediction

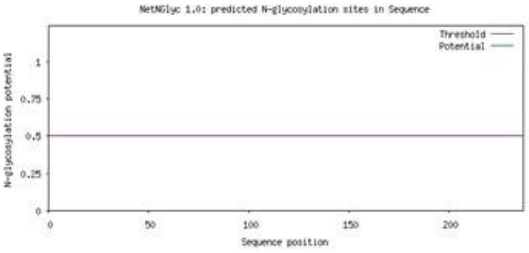


Myx1UPO

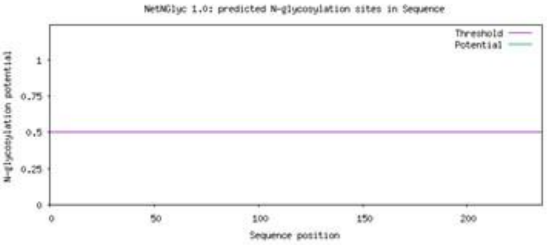


Myx2UPO

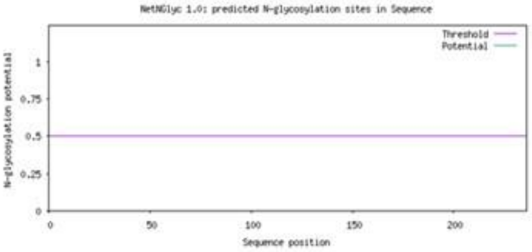


Del1UPO

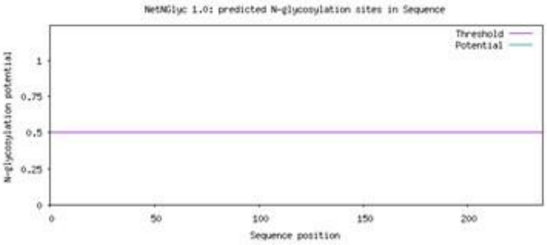


Del2UPO

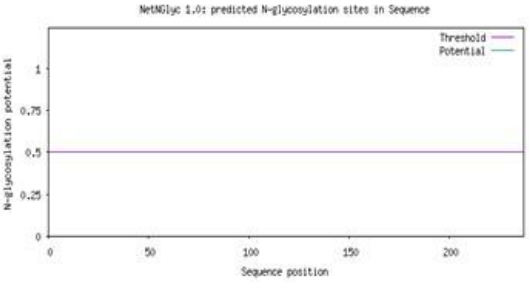


PolUPO

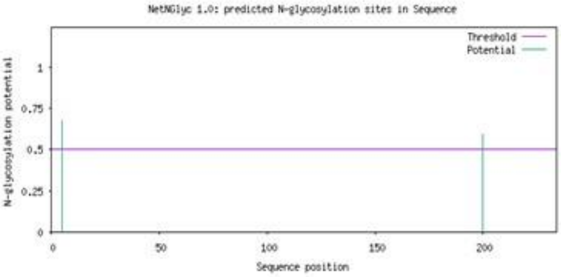


PseUPO

| SeqName | Position | Potential | N-Glyc |
|---|---|---|---|
| Sequence | 5 NATD | 0.6789 | + |
| Sequence | 200 NITL | 0.5949 | + |

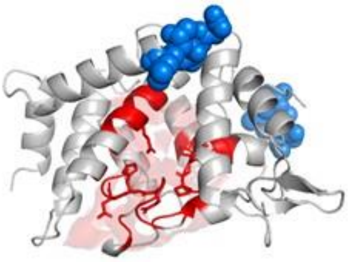

**Figure S6.** Glycosylation site analysis

PseUPO contains two potential glycosylation sites. Spatial structure analysis (with the active site as the reference) revealed that these sites are located 5 Å away from the active center, indicating that the glycosylation sites are distant from the active region. This spatial distribution characteristic suggests that the glycosylation modification of PseUPO (marked in blue in the schematic diagram) does not significantly affect its catalytic activity.

**Table S1.** Basic properties of the enzymes

| | Myx1UPO | Myx2UPO | PseUPO |
|---|---|---|---|
| Molecular Formula | $C_{1070}H_{1760}N_{326}O_{339}S_6$ | $C_{1071}H_{1771}N_{327}O_{333}S_8$ | $C_{1059}H_{1727}N_{327}O_{331}S_5$ |
| Molecular Weight (Da) | 24808.08 | 24813.31 | 24496.65 |
| Theoretical pI | 7.79 | 7.75 | 7.16 |
| Instability Index | 37.35 | 37.61 | 35.59 |
| Grand Average of Hydropathicity (GRAVY) | -0.288 | -0.216 | -0.243 |
| Molar Extinction Coefficient ($mL \cdot mg^{-1} \cdot cm^{-1}$) | 0.3 | 0.3 | 0.3 |
| Most Hydrophobic Position | 213 | 170 | 209 |
| Hydrophobicity at Maximum Position | 1.833 | 2.178 | 2.233 |
| Most Hydrophilic Position | 16 | 16 | 229 |
| Hydrophilicity at Maximum Position | -2.256 | -2.256 | -2.367 |

**Table S2.** Basic properties of the enzymes

| | Del1UPO | Del2UPO | Del2UPO |
|---|---|---|---|
| Molecular Formula | $C_{1068}H_{1766}N_{326}O_{332}S_7$ | $C_{1069}H_{1766}N_{326}O_{334}S_7$ | $C_{1083}H_{1785}N_{333}O_{340}S_8$ |
| Molecular Weight (Da) | 24710.17 | 24754.18 | 25167.59 |
| Theoretical pI | 7.76 | 7.09 | 8.47 |
| Instability Index | 36.07 | 35.43 | 38.47 |
| Grand Average of Hydropathicity (GRAVY) | -0.174 | -0.197 | -0.326 |

| | | | |
|---|---|---|---|
| Molar Extinction Coefficient ($mL\cdot mg^{-1}\cdot cm^{-1}$) | 0.3 | 0.3 | 0.29 |
| Most Hydrophobic Position | 170 | 170 | 120 |
| Hydrophobicity at Maximum Position | 2.178 | 2.178 | 1.489 |
| Most Hydrophilic Position | 16 | 16 | 16 |
| Hydrophilicity at Maximum Position | -2.256 | -2.256 | -2.256 |

**Table S2.** UPO enzyme activity and catalytic efficiency

| **UPO** | **NBD (Dealkylation → Catechol)** | | **Veratryl Alcohol (Hydroxyl Oxidation → Veratraldehyde)** | | **Naphthalene (Hydroxylation → 1-Naphthol)** | |
|---|---|---|---|---|---|---|
| | Catalytic Activity （U/L） | kcat/Km （$mM^{-1}\cdot s^{-1}$） | Catalytic Activity （U/L） | kcat/Km （$mM^{-1}\cdot s^{-1}$） | Catalytic Activity （U/L） | kcat/Km （$mM^{-1}\cdot s^{-1}$） |
| Myx1UPO | 1147.00 | 78.95 | 978.00 | 248.59 | 1959.00 | 1063.58 |
| Myx2UPO | 462.50 | 86.21 | 270.77 | 109.77 | 541.74 | 1033.53 |
| PseUPO | 703.68 | 89.40 | 609.33 | 118.38 | 1219.26 | 1716.79 |
| Del1UPO | 319.50 | 14.93 | 300.35 | 73.28 | 600.98 | 944.89 |
| Del2UPO | 271.16 | 13.12 | 254.95 | 94.79 | 510.04 | 1441.20 |
| PolUPO | 663.00 | 77.64 | 584.24 | 120.71 | 1149.11 | 1397.74 |

**Amino acid sequences of the enzymes：**

Myx1UPO is derived from Myxococcales bacterium (GenBank accession number: MBL8634636.1)：

MSKQSEPAASGWQPAQPGEQRSPCPALNSLANGGDIPRSGIATRAQLVSAMVNRLGVST

ALADKLAKAALEQFGKPGEGSEPVLHLQDLCQHGKLEHDASLTRQDAHAGDNAKVDPA

LIEQLLSLSKNGQTLTLDDLATAHQVRMQQSAQGGHQVPSKAGFVGTVEASLLYNVLSR

GPQGISLADAREFLLHERVPEGLSGRGISIGQVAANAITIAIKGNLPLCESARRAKDALKK

Myx2UPO is derived from Myxococcales bacterium (GenBank accession number: MBL9044159.1)：

MSQHSQAGASDWQPAQPGEQRSPCPALNSLANGGHIPRSGIASRAQLVSAMVQRLGLAT
ALADKLAKEAIEKFGKPGDGGEQVLHLQDLCQPGKLEHDASLTRQDAHVGDSAKIDPAL
VEQLLALSKNGQTLTLDDLATAHQIRMQQSAQGGHVVPPKAGLVGTVEAALLYCVLSR
GEGISLADAREFLLSERIPQGLTGHDISLAQVARRAITVALKGNLPMCDAARRAKDALKK

PseUPO is derived from Pseudomonadota bacterium (GenBank accession number: HND09080.1)：

MSQNNATDWQPAQPGEGRSPCPALNSLANGGHLPRNGIASRSQLITAIVGQLGISTEFANT
LANAALEKFGKPGDGGEQVLHLQDLCQHGKLEHDASLTRQDAHAGDHAKVDPSLVEQL
LSLSKNGQTLTLDDLAAAHQIRMHQSAQGGHHVPGKAGFVGTVEAALLYTVLARGESGI
SLADAREFLLTERVPQGLAGRNITLAQVAVRAIAIAAKGNLPVCEAARRAKEATKK

Del1UPO is derived from Deltaproteobacteria bacterium (GenBank accession number: MBP6609162.1)：

MSQHTQAGASEWQPAQPGEQRSPCPALNSLANGGHIPRSGIASRAQLVSAMVQRLGLAT
ALADKLATEALAKFGKPGDGGEQVLHLQDLCQPGKLEHDASLTRQDAHVGDSAKIDPA
LVEQLLALSKNGQTLTLDDLATAHQIRMQQSAQGGHVVPPKAGLVGTVEAALLYCVLSR
GEGISLADAREFLLSERIPQGLTGHDISLAQVARRAVTVALKGNLPVCDAARRAKDALKK

Del2UPO is derived from Deltaproteobacteria bacterium (GenBank accession number: MBP8195526.1)：

MSQHTQAGASEWQPAQPGEQRSPCPALNSLANGGHIPRSGIASRAQLVSAMVQRLGLAT
ALADKLATEALEKFGKPGDGGEQVLHLQDLCQPGKLEHDASLTRQDAHVGDSAKVDPA
LVEQLLALSKNGQTLTLDDLATAHQIRMQQSAQGGHVVPPKAGLVGTVEAALLYCVLSR
GEGISLADAREFLLSERIPQGLTGHDISLAQVARRAVTVALKGNLPVCDAARRAKDALKK

PolUPO is derived from Polyangia bacterium (GenBank accession number: MFO0622164.1)：

MSKQSEPAASGWQPAQPGEQRSPCPALNSLANGGHIPRSGIATRAQLVSAMVNRLGVST
ALADKLAVAAMEKFGKPGEGSEQVLHLQDLCEHGKLEHDASLTRQDTHVGDNAKVDP
ALVEQLLSLSKNGQTLTLDDLATAHQIRMQQSAQGGHQVPSKAGFVGTVEASLLYNVLS
RGQNGISLADAREFLLHERIPEGLTGRDISMPQVARRAITIALKGNLPICDAARRAKDALK
K

**Determination of UPO Enzymatic Activity towards Aromatic Substrates**

The maximum characteristic absorption wavelengths of the products (4-nitrocatechol, veratraldehyde, and 1-naphthol) were 425 nm, 310 nm, and 324 nm, respectively. Standard concentration curves were constructed by plotting absorbance against corresponding product concentrations, and the respective linear regression equations were derived for calculating product concentrations.

Preparation of Standard Solutions and Standard Curves Standard solutions of the three products were prepared at the following concentrations:

4-nitrocatechol: 0.05, 0.1, 0.2, 0.3, 0.4, 0.5, 0.6, 0.7, 0.8, 0.9, and 1.0 mmol/L;

Veratraldehyde: 0.5, 0.8, 1.0, and 2.0 mmol/L;

1-naphthol: 0.8, 1.0, 2.0, 4.0, and 8.0 mmol/L.

For each standard sample, the reaction mixture (total volume: 200 μL) was composed of 30 μL of the standard solution, 1 μL of diluted $H_2O_2$ (30% $H_2O_2$ diluted 10-fold), 30 μL of ultrapure water, and 139 μL of pH 7.0 PBS buffer. The standard curves are shown in Figure 8, with the linear regression equations as follows (y: absorbance; x: product concentration, mmol/L):

4-nitrocatechol:

y=0.8788x+0.0973

Veratraldehyde:

y=0.5839x+0.1298

1-naphthol:

y=0.1546x+0.0768

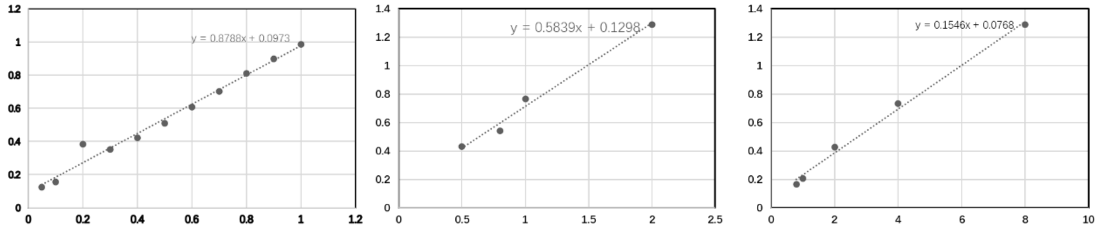


A B C

Figure S6. Standard concentration curve: A 4-nitrocatechol; B: Fenucraldehyde C: 1-naphthol

**Enzymatic Reactions and Activity Calculation towards Aromatic Substrates**

After establishing the standard curves, the enzymatic catalytic reactions of the substrates were performed as follows:

In a 2 mL centrifuge tube, 30 μg of purified UPO protein was mixed with 30 μL of substrate solution (concentration: 6.6 mmol/L; NBD and resveratrol were dissolved in acetonitrile, while naphthalene was dissolved in ethyl acetate).

One microliter of diluted $H_2O_2$ (30% $H_2O_2$ diluted 10-fold) was added, and the total volume was adjusted to 200 μL with cell lysis buffer.

The 2 mL centrifuge tube was incubated in a metal bath at 25 °C with shaking at 200 rpm for 5 min. After incubation, the mixture was centrifuged at 12,000 rpm for 2 min, and the supernatant was transferred to a 96-well plate.

The absorbance of the supernatant was measured using a microplate reader at the respective characteristic wavelengths of the products.

Product concentrations were calculated using the aforementioned linear regression equations. Enzymatic activity (expressed as $mmol \cdot min^{-1} \cdot L^{-1}$, equivalent to 1000 U/L) was further calculated as the ratio of product concentration (mM) to reaction time (min).

**Determination of In Vitro Catalytic Kinetic Parameters towards Aromatic Substrates**

The kinetic parameters were determined based on the enzymatic activity assay method described in Example 3, with the following specific procedures:

1. Determination of Maximum Reaction Rate (Vmax) and Turnover Number (kcat)

For each UPO, the maximum reaction rate (Vmax) was measured by monitoring the absorbance at 425 nm ($A_{425}$) at 2-minute intervals throughout the enzymatic reaction. The product concentration at each time point was calculated using the pre-established standard curve (Figure S6), and Vmax was derived from the linear phase of the reaction progress curve. The turnover number (kcat) was calculated according to the molar amount of the added UPO enzyme (kcat = Vmax / $[E]_0$, where $[E]_0$ represents the initial enzyme concentration).

2. Determination of Michaelis-Menten Constant (KM)

NBD was used as the substrate to determine the Michaelis-Menten constant (KM) of each UPO. The reaction system was configured as follows:

Substrate concentration gradient: 0.1–6.6 mM (30 μL per reaction);

Enzyme dosage: 30 μg of purified UPO;

Reaction conditions: Incubated at 25 ℃ for 5 min;

Reaction termination: 20 μL of 30% (v/v) HCl was added to quench the reaction;

Post-reaction processing: The mixture was centrifuged, and the supernatant was transferred to a 96-well plate. The absorbance at 425 nm was measured using a microplate reader, and the product concentration was calculated based on the standard curve (Figure S6).

The Michaelis-Menten constant (KM) for each enzyme was obtained by fitting the concentration-response data to the Michaelis-Menten equation using GraphPad Prism software. The results are summarized in Table S2.

**Catalytic Reaction System for Terpenoid Substrates:**

The total volume of the reaction system was 2.5 mL, with the following specific formulation: Acetone (as a cosolvent) accounting for 16% of the total volume was added to the system to achieve a final substrate concentration of 10 mM, and the remaining volume was made up with pH 7.0 buffer. At the initial stage of the reaction, 160 μL of enzyme solution and 10 μL of 3% (mass concentration) $H_2O_2$ were added to the aforementioned system. During the reaction, 30 μL of enzyme solution and 9 μL of 3% (mass concentration) $H_2O_2$ were supplemented every 20 minutes. The reaction system was incubated in a constant-temperature metal bath at 25 ℃ with shaking at 600 r/min for a total reaction time of 2 hours. After the reaction, 400 μL of ethyl acetate was added for extraction. Subsequently, 175 μL of the extract was mixed with 25 μL of internal standard solution (dodecane, concentration: 10 mM) and prepared for detection.

**GC-MS Detection Method and Product Selectivity Calculation:**

Chromatographic conditions: Column: HP-5ms capillary column; Ion source: Electron ionization (EI), 230 eV; Injection volume: 1 μL; Injection temperature: 250 ℃; Detector temperature: 325 ℃; Column temperature program: Initial temperature of 80 ℃ (held for 5 min), followed by heating to 160 ℃ at a rate of 20 ℃/min and maintaining for 18 min.

Quantification method: The product yield was determined using the internal standard method with dodecane as the internal standard. A 50 mg/L standard solution was prepared, and 175 μL of the sample was mixed with 25 μL of 50 mg/L dodecane to obtain a final dodecane concentration of 5 mg/L. The sample yield was calculated based on the peak area ratio of the product to the standard.
